# Bone Marrow-Derived Cells Fuse with Bergmann Glia in Experimental Autoimmune Encephalomyelitis

**DOI:** 10.64898/2026.09.29.755471

**Authors:** Pablo G. Téllez de Meneses, Sara Sánchez-Monreal, Carmelo A. Ávila-Zarza, Carla Escudero-Solano, Eduardo Weruaga, David Díaz, Jorge Valero, José Ramón Alonso

**Author notes:** Equal contribution (Jorge Valero, José Ramón Alonso).

## Abstract

Bone marrow-derived cells (BMDCs) can infiltrate the brain and adopt new identities through transdifferentiation or fusion, yet their interactions with specific neural cells remain poorly defined. We examined BMDC fusion with Bergmann glia (BG), a key cerebellar support cell, in a mouse model of multiple sclerosis. After bone marrow transplantation, BMDCs fused with BG to form heterokaryons, retaining BG morphology and molecular markers. These heterokaryons contained donor and host nuclei that differed in both morphology and transcriptional activity. Remarkably, the BMDC nucleus localized not only to the soma but also along BG processes. This discovery identifies BMDC-BG fusion as a previously unrecognized form of glial plasticity, opening avenues for therapeutic strategies aimed at potentiating glial-mediated repair.

## INTRODUCTION

The fusion of bone marrow-derived cells (BMDCs) with Purkinje neurons was discovered in the early 2000s in both mice and humans ^1,2^. This fusion leads to a singular Purkinje cell called heterokaryon, in which both the original neuronal and BMDC nuclei are preserved and transcriptionally active ^3,4^.

Fusion events have been observed under physiological conditions ^5^, but the formation of stable heterokaryons has only been described after bone marrow transplantation ^3,4^. The number of heterokaryons is generally low, but it increases with age and after irradiation or partial ablation of Purkinje cells ^6–9^. In addition, BMDC fusion also increases in specific pathologies, such as cerebellar ataxia ^9–11^ and, especially, in multiple sclerosis, due to autoimmunity and chronic inflammation ^4,12,13^.

The observation that cell fusion events increase in neurodegenerative contexts has motivated intense investigation into their therapeutic potential for brain repair, as a means to rescue or protect degenerating cells that may lack the expression of significant genes ^13–18^. However, the scarcity of heterokaryon Purkinje cells hinders detailed analysis of fusion mechanisms ^19–21^. Expanding our understanding of cell fusion beyond Purkinje cells may overcome this limitation and reveal the general mechanisms governing cell fusion under physiological conditions or during neurodegeneration.

Early studies suggested that fusion within the nervous system was restricted to Purkinje cells ^4,22^. Nevertheless, subsequent studies have identified fusion events with different neuronal types in pathological contexts ^23,24^. Finally, the interest in studying fusion events led to the discovery that fusion can happen not only with neurons but also with pericytes ^25^, astrocytes from the substantia nigra pars compacta ^24^, and Müller glial cells ^26^, revealing cell fusion as a more prevalent phenomenon across the nervous system than previously anticipated. However, fusion with other glial types remains largely unexplored, representing a significant gap in our understanding.

To address this, here we focus on the study of Bergmann glia (BG). BG are specialized radial astrocytes that extend processes along the cerebellar cortex, physically interacting with Purkinje cell somata and dendrites ^27^. During cerebellar development, they serve as a migratory scaffold that guides granule cells from the external to internal granular layer, providing essential cellular interactions for proper lamination and circuit assembly ^28^. Moreover, BG have been related to survival, dendritic growth, and spine formation in Purkinje cells ^29,30^. They express characteristic astrocytic markers, such as the glial fibrillary acidic protein (GFAP) ^31^ and S-100β ^32^, and stem cell markers, including Sox1, Sox2, and Sox9 ^33^, suggesting new roles for them as stem cells in the mature cerebellum ^34^. Even though Chen et al. (2011) ^15^ mentioned -but did not show- the existence of GFP^+^ putative BG after GFP^+^ BMDC transplantation, they did not explore if they originated by fusion or other mechanisms such as “transdifferentiation”^35^.

Here, through exhaustive immunofluorescence analysis combined with high-resolution microscopy and advanced image quantification, we provide evidence for the first time that BMDCs can fuse with BG in the cerebellum during Experimental Autoimmune Encephalomyelitis (EAE, an experimental model of multiple sclerosis), forming heterokaryons. These fused cells preserve the characteristic morphology and molecular markers of BG, while harboring two distinct nuclei, consistent with a fusion event. This previously unrecognized phenomenon refines our understanding of BMDC plasticity and supports the development of stem cell–based therapeutic strategies targeting cerebellar neurodegeneration and, more broadly, glial dysfunction.

## MATERIALS AND METHODS

### Animals

Twelve BALB/c × C57BL/6J × DBA/2J congenic mice ^18^ (Mus musculus, L. 1758; Jackson Laboratories), between two and five months old, were used as bone marrow donors. These animals exhibited constitutive green fluorescent protein (GFP) expression, enabling easy visualization of their cells in recipients. Forty C57BL/6J × DBA/2J congenic wild-type mice were used as recipients. These animals were handled in accordance with the European Union directive (2010/63/EU) and the Spanish Current Law (RD 118/2021) on the maintenance, use, and care of laboratory animals. All the experimental procedures for this study were approved by the Bioethics Committee of the Universidad de Salamanca and the regional government of Castilla y León (Salamanca, Spain, Refs 613 and 1258).

### Bone marrow isolation and BMDC transplantation

Donor mice were sacrificed by cervical dislocation and decapitation. Femurs, tibias, and hips were dissected, and the bone marrow was extracted using a 30G insulin syringe and α-MEM no nucleoside medium (Gibco, 22561054). Bone marrow was passed through a 70 μm cell strainer (Falcon, 352350), and isolated cells were collected in 50 mL sterile tubes (VWR, 525-1107) and suspended in α-MEM no nucleoside medium. Cells were centrifuged (4 °C, 5 min, 1500 rpm) and then resuspended in 5 mL of lysis buffer (0.83% w/v NH4Cl, 0.1% w/v KHCO3, 0.372% w/v EDTA, pH 7.3) to lyse erythrocytes. After 5 minutes at room temperature, Dulbecco’s Phosphate-Buffered Saline (DPBS; Gibco, 14190144) was added to a final volume of 50 mL to stop the lysis buffer reaction. At this point, 50 μL were collected from the center of the tube and counted using a Neubauer chamber to estimate the total number of gathered cells. While counting, cells were centrifuged again as previously described and resuspended in DPBS to a final concentration of 7.5 x 106 cells per 50 μL, which was the volume used for the transplant.

Recipient animals were exposed to a single whole-body irradiation with a minimal lethal dose of 7.5 Gy at the age of 3 weeks ^36^. Irradiation was performed using a 137Cs gamma irradiator, Gammacell 1000 Elite (MDS Nordion). 24 hours after irradiation, animals were transplanted retro-orbitally with 50 μL of DPBS containing 7.5 x 106 cells. For the transplant, mice were anesthetized with inhalational isoflurane (induction: 4% v/v in oxygen at 0.8 L/min; maintenance: 2% v/v in oxygen at 0.8 L/min) delivered via a precision vaporizer using an induction chamber and a nose cone for maintenance. Transplanted animals were housed separately from other animals and monitored to ensure full recovery.

### EAE induction

Approximately one month after the transplant, EAE was induced in recipient animals. Animals were anesthetized as previously described and received four subcutaneous injections in the flanks. A 1:1 emulsion of myelin oligodendrocyte glycoprotein peptide (MOG_35-55_; enQuireBio, 15981319) in filtered 0.1 M phosphate-buffered saline (PBS; pH 7.4) and Complete Freund’s Adjuvant (CFA) was prepared. A total volume of 200 μL containing 75 μg (n = 8), 100 μg (n = 8), or 200 μg (n = 14) of MOG was administered per animal. The CFA was prepared from Incomplete Freund’s Adjuvant (Thermo Scientific 77145) supplemented with inactivated Mycobacterium tuberculosis (BD 231141) at a concentration of 2 mg/mL. Just before the emulsion injection and 48 hours later, animals were intraperitoneally injected with 200 μL of filtered PBS containing 200 ng of pertussis toxin (Tocris 3097). Control animals (n = 10) underwent identical procedures with ovalbumin (OVA) as a non-encephalitogenic peptide substitute for MOG^35–55^.

Mice were monitored daily for body weight and clinical signs from EAE induction through peak disease. Following clinical stabilization or complete symptom resolution, monitoring continued less frequently, depending on the animal’s condition, until the study endpoint. Clinical severity was evaluated using a standardized clinical scoring scale ranging from 0 to 5 (0, no symptoms; 0.5, tail weakness; 1, tail paralysis; 1.5, unilateral hind limb paresis; 2, bilateral hind limb paresis; 2.5, unilateral hind limb paralysis; 3, bilateral hind limb paralysis; 3.5, unilateral forelimb paresis; 4 bilateral forelimb paresis; 4.5 unilateral forelimb paralysis; 5, bilateral forelimb paralysis, moribund). Animals reaching a score ≥3.5 were scheduled for euthanasia per predetermined humane endpoints; however, no animal exceeded a maximum score of 3. To compare symptomatology across animals, we used the cumulative disease index (CDI; ^37–39^), which is the sum of each animal’s daily clinical score. Because the duration of clinical follow-up differed among animals, CDI values were normalized by dividing them by the number of days for which clinical scores were recorded.

### Immunofluorescence

Animals were sacrificed either eight or fourteen weeks after the EAE induction. Under deep general anesthesia, animals were injected with heparin in the left heart ventricle to avoid clot formation. Saline solution 0.9% was passed from the left ventricle along the circulatory system, and the right atrium was opened to allow blood flow. Once the circulatory system was rinsed, Somogyi’s fixative (4% w/v paraformaldehyde and 15% v/v saturated picric acid in 0.2 M phosphate buffer, pH 7.4, PB) was perfused for 15 minutes. After the fixation, the brain was dissected out, cut into pieces (hemispheres and two halves of the cerebellum), and postfixed in Somogyi’s fixative for 2 hours. Finally, pieces were rinsed two times with 0.1M PB and cryoprotected in 30% w/v sucrose in 0.1M PB until they sank. Finally, samples were frozen at -80 °C for storage.

Sagittal sections of the cerebellum (40 μm-thick) were obtained using a freezing sliding microtome (Leica Frigomobil, Jung SM 2000). Slices were then rinsed in 0.1M PB and stored in an antifreezing solution (30% v/v glycerol, 30% v/v polyethylene glycol in 0.1M PB) when the immunofluorescence technique was not performed within the following days.

One section from every 6 consecutive sections from the half cerebellum was selected for immunofluorescence analysis. Slices were rinsed in PBS (3 x 10 min) and then incubated with the primary antibody solution for 24-72 hours at 4 °C. This solution was composed of 5% v/v donkey serum (Linus S02171), 0.2% v/v Triton X-100 (Probus 22.7810), primary antibodies (Table 1), and PBS. After incubation, slices were rinsed in PBS (3 x 10 min) and incubated in the secondary antibody solution for 1 hour (2 h for the S100β antibody to enhance tissue penetration) at room temperature. For stimulated emission depletion (STED) microscopy, the sample was incubated in the secondary antibody overnight at 4 °C. The composition of this solution was the same as for the primary antibody solution, but with secondary (Table 1) instead of primary antibodies, and it also included DAPI (4′,6-Diamidine-2′-phenylindole dihydrochloride, Sigma- Aldrich D9542) at a final concentration of 0.1 μg/mL. After the incubation with this solution, slices were rinsed in PBS (3x10 min), mounted on gelatinized slices, and covered with a homemade antifading solution (70% v/v glycerin, 5% w/v N-propyl gallate, 0.51% w/v NaCl, 0.42% w/v glycine, 0.021% w/v NaOH, and 0.03% w/v sodium azide) and coverslips (VWR ECN 631-1574).

### Data collection and image analysis

GFP^+^ BG (derived from transplanted bone marrow) cells were counted using an epifluorescence microscope (Olympus Provis AX70) equipped with a UPlanSApo 40x/0.90 dry objective. Cell locations were marked for each animal on printed mosaic images of the DAPI staining acquired with a Leica Stellaris DMi8 laser confocal microscope (Leica Microsystems) using a HC PL FLUOTAR 4x/0.13 DRY objective. Then, cells from different animals were represented in a cerebellum scheme to facilitate visualization. All vermal sections were combined into a single schematic overview. In contrast, non-vermal sections were arranged into two centro-lateral schemes according to the presence or absence of the pons and most of the medulla. Finally, cell distribution was compared between the vermal and non-vermal regions. Within the vermis, cell distribution was analyzed among the different domains ^40^ and lobes.

Mosaics and images for colocalization and nuclei analysis were obtained using a Leica Stellaris DMI8 laser confocal microscope and HC PL APO CS2 20x/0.75 DRY or HC PL APO CS2 40x/1.30 OIL objectives. Cell reconstruction in Fig. 2C (purple) was generated by manually tracing all processes of a single BG cell, beginning at the soma optical section and following each process across successive planes.

**Figure 1.**
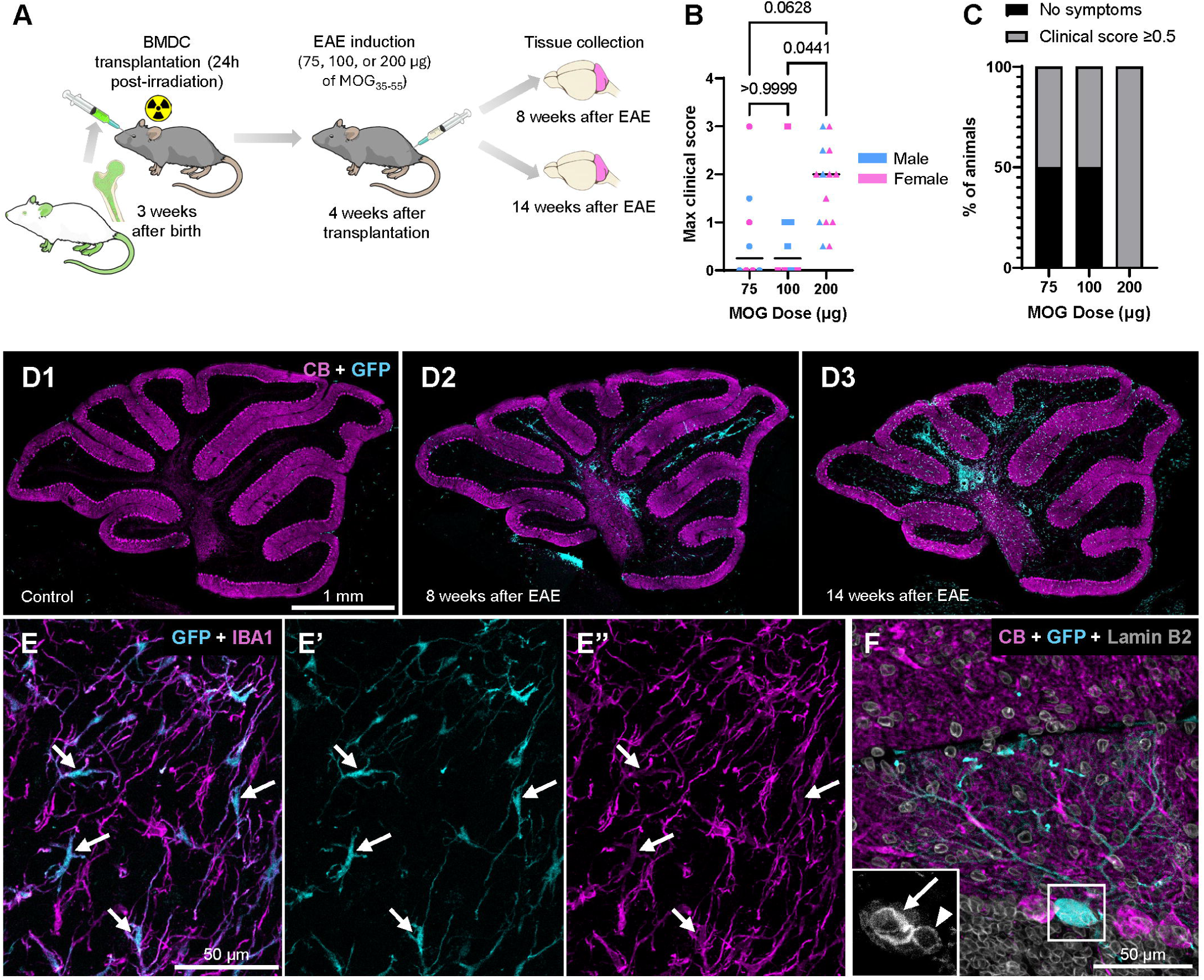
BMDC infiltration and fusion in the cerebellum after EAE induction. (A) Schematic representation of the experimental design used to assess bone marrow– derived cell (BMDC) infiltration and fusion in the EAE mouse model. Illustrations were adapted from NIH Bioart source (https://bioart.niaid.nih.gov) using Inkscape. (B) Maximum clinical scores reached for the different doses of MOG_35-55_ used to induce EAE. (C) Percentage of animals remaining asymptomatic across the doses of MOG_35-55_ tested (75 μg, n = 8; 100 μg, n = 8; 200 μg, n = 14) (D) Maximum-intensity projections of confocal mosaic images of cerebellar vermis sections showing Purkinje cells labeled with calbindin (magenta) and GFP^+^ infiltrated BMDCs (cyan) in a control mouse (ovalbumin induction; D1), and in mice sacrificed 8 (D2) and 14 (D3) weeks after EAE induction. (E) Optical section of GFP^+^ infiltrated cells (cyan; E’) showing colocalization with the microglial/macrophage marker Iba1 (arrows; magenta; E’’). (F) Optical section of a fused GFP⁺ Purkinje cell expressing the Purkinje-specific marker calbindin. The white box demarcates the soma of the heterokaryon Purkinje cell. The inset displays the isolated Lamin B2 channel (gray) of the boxed area, revealing two distinct nuclei (arrow and arrowhead) sharing the same cytoplasm. CB, calbindin.

**Figure 2.**
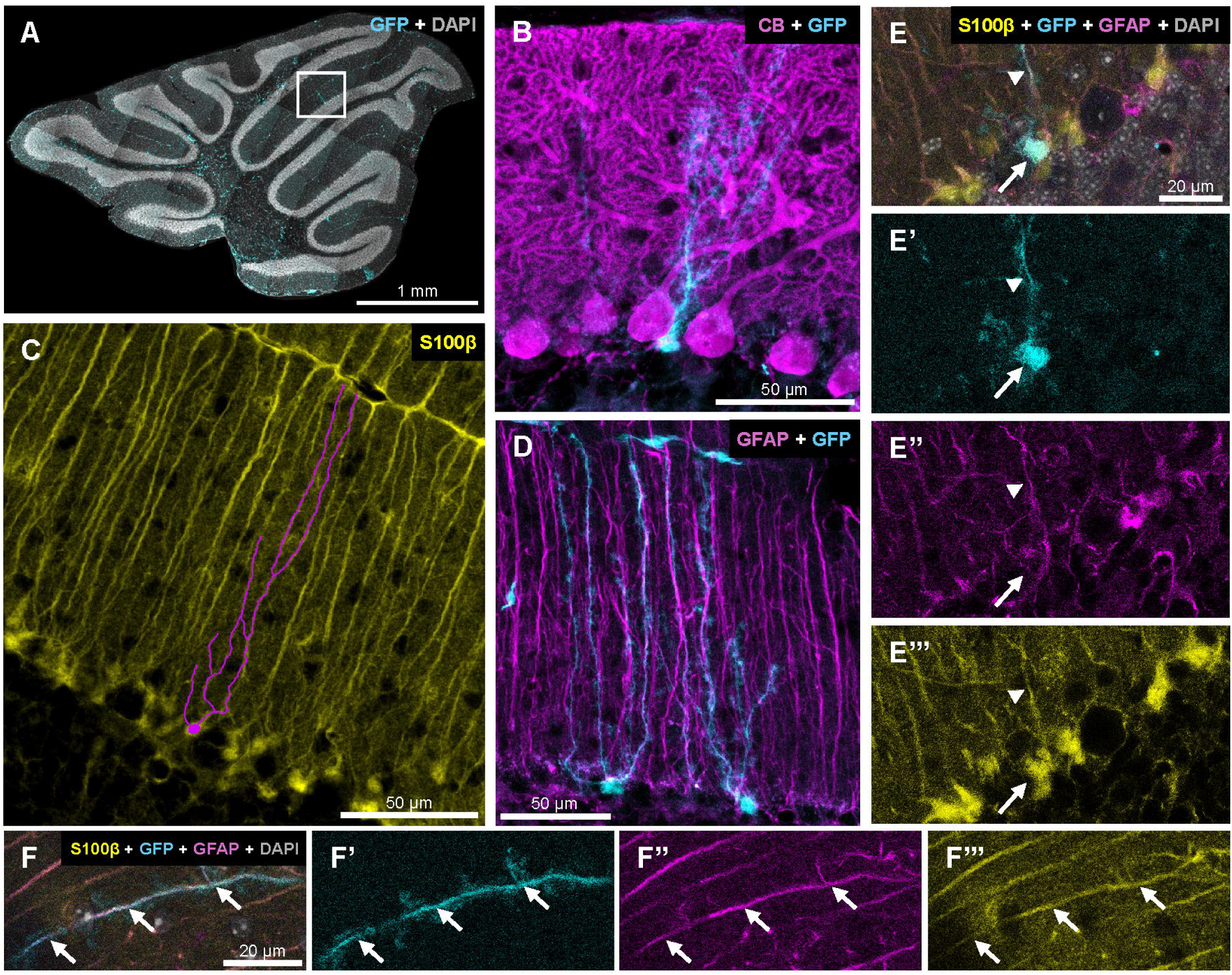
Identification of GFP^+^ cells with a BG phenotype. (A) Maximum-intensity projection of a confocal mosaic image of the cerebellar vermis showing DAPI (gray) and GFP^+^ cells (cyan). The white box indicates the location of a GFP^+^ BG cell. (B) Maximum-intensity projection of a GFP^+^ BG cell located among Purkinje cells, labeled with calbindin (magenta). (C) Maximum-intensity projection of S100β immunostaining (yellow) showing multiple BG cells, with one reconstructed cell in magenta. (D) Maximum-intensity projection of two GFP^+^ BG cells (cyan) displaying typical GFAP staining (magenta) and the characteristic BG morphology. (E) Single optical plane showing a GFP^+^ BG cell (cyan; E’) soma with GFAP (magenta; E’’) and S100β (yellow; E’’’) colocalization. (F) Single optical section showing a GFP^+^ BG cell process (cyan; F’) with GFAP (magenta; F’’) and S100β (yellow; F’’’) colocalization. CB, calbindin.

FIJI software ^41^ was used for image analysis. Nuclei measurements were performed by creating regions of interest (ROIs) in z-stack images and selecting the optical section where the area of the nucleus was largest. Within these ROIs, we measure eight different parameters (area, perimeter, width, height, major and minor axis lengths of the best fitting ellipse, Feret’s diameter or maximum caliper, and minimum Feret’s diameter or minimum caliper) for Principal Component Analysis (PCA) and the roundness (Equation 1) for morphological analysis.

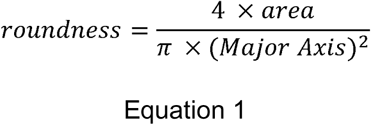

A subset of the previous ROIs was also used to analyze DAPI and the active form of RNA polymerase II staining using the Thresholderer FIJI macro (https://github.com/Jorvalgl/Thresholderer). The macro quantifies pixel coverage area as a percentage, mean intensity, and integrated density across sequential intensity thresholds (each threshold range spanning from the threshold value to the maximum intensity value of 255). A rolling-ball background subtraction was applied to all images (radius 50 pixels for DAPI and 100 pixels for RNA Pol II) to correct for uneven background. The mean percentage of area covered for thresholds ranging from 0 to 150 was plotted for both markers to evaluate differences between BMDC and BG nuclei. The DAPI channel was then thresholded at an intensity value of 40 to selectively segment heterochromatin-enriched regions, whereas the RNA Pol II channel was thresholded at an intensity value of 30 to capture all detectable active transcriptional foci.

Finally, the distance between nuclei was determined by obtaining centroid coordinates (x, y, z) for each nuclear ROI pair and calculating three-dimensional Euclidean distance using Equation 2.

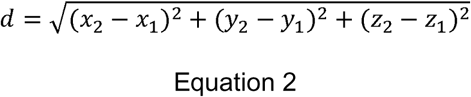

A GFP^+^ cell was segmented for Fig. 4A and supplementary video 1. The original dataset consisted of a multichannel z-stack image. A Gaussian blur filter (radius = 2) was first applied to all z-planes and channels to reduce noise. Segmentation of GFP⁺ structures was performed on the GFP channel using the Labkit plugin ^42^. The resulting binary mask of the cell of interest was generated using the 3D Object Counter tool. To isolate the corresponding regions in the other channels, the Image Calculator tool was used with the “AND” operator to combine the mask with the GFP, DAPI, and GFAP channels individually. These three processed channels were subsequently merged using the “Merge Channels” option to reconstruct the multichannel image of the selected GFP⁺ cell. The final image thus displays only structures corresponding to the GFP⁺ cell across all channels. The same method with channel variations was used for the Purkinje cell soma detail. 3D reconstructions and supplementary video 2 were made using LasX software.

**Figure 3.**
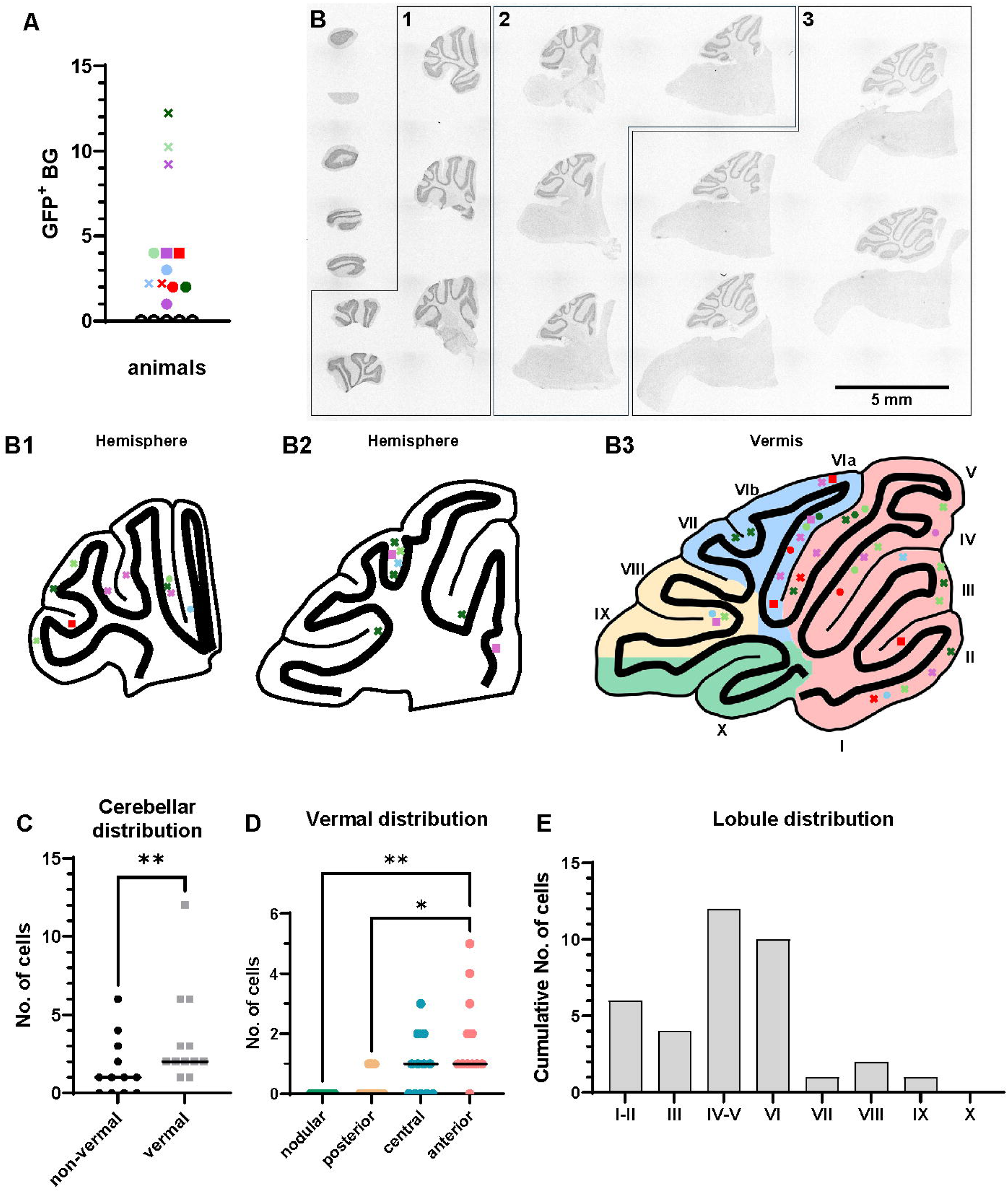
Topographic mapping of GFP^⁺^ Bergmann glia in the cerebellar cortex. (A) Scatter plot showing the total number of GFP^+^ Bergman glia (BG) identified in 2 cerebellar series per animal (n = 17). Each specific color/symbol combination represents data from a different animal. White dots indicate animals in which no GFP⁺ BG cells were detected: 3 controls and 2 EAE mice. (B) Representative mosaic image of a complete cerebellar series labeled with DAPI (gray), illustrating the anatomical grouping of sections (B1–B3, flocculus was discarded for quantifications). Colored symbols map the location of GFP⁺ BG cells for individual animals, maintaining the coding from (A). (C) Analysis of the cerebellar distribution of GFP^+^ BG cells; each dot/square represents the number of cells in each animal. Solid line indicates the median (n = 12; Wilcoxon test). (D) Vermal distribution of the total number of GFP^+^ BG cells (corresponding to B3 grouping and colors of regions); each dot represents the number of cells in each animal. Solid line indicates the median (n = 12; Friedman test with Dunn’s multiple comparisons test). (E) Frequency distribution of GFP^+^ BG cells across cerebellar lobes (see B3 grouping). Bars represent the total number of cells pooled across all animals (n = 12). * p < 0.05; ** p < 0.01.

**Figure 4.**
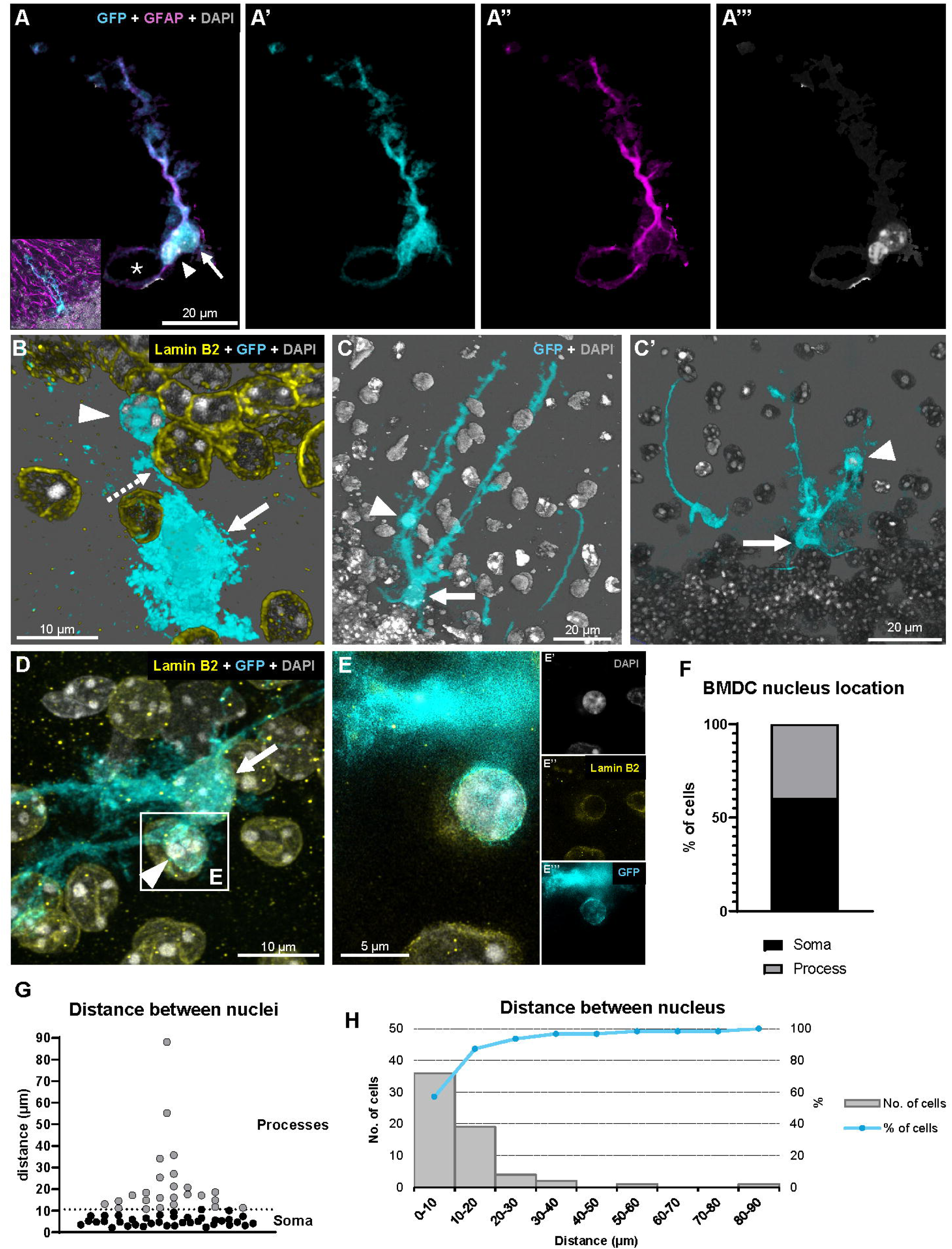
Fusion of BG with BMDCs revealed by three-dimensional imaging and nuclear positioning analysis. (A) Single optical section reconstruction of a fused BG, in which two different nuclei stained with DAPI (gray; A’’’; arrow and arrowhead) can be distinguished, both surrounded by GFP (cyan; A’) and GFAP (magenta; A’’) staining. Note the characteristic BG process enwrapping the unstained Purkinje cell somata (asterisk). (B) Three-dimensional reconstruction from a confocal z-stack of a fused GFP^+^ BG with an extrasomatic nucleus (arrowhead) displaced toward the granule cell layer. Nuclei are labeled with DAPI and Lamin B2 (yellow). The process connecting the somatic (arrow) and extrasomatic (arrowhead) nuclei is pointed by a dotted arrow. (C- C’) Different views of a three-dimensional reconstruction from a confocal z-stack of a fused BG with an extrasomatic nucleus displaced towards the molecular layer (arrowhead), fully enclosed by GFP staining; the arrow points to the somatic nucleus. (D) Single optical section of a fused GFP^+^ BG cell with an extrasomatic nucleus towards the molecular layer (arrowhead), detailed in E; the arrow points to the somatic nucleus. (E) STED super-resolution image of the boxed region in (D), showing that the nucleus labeled with DAPI (grey; E’) and Lamin B2 (yellow; E’’) is contained within the GFP⁺ BG cell (cyan; E’’’). (F) Quantification of nuclear positioning: percentage of fused BG cells in which the BMDC nucleus is in the soma versus in a process (n = 62). (G) Analysis of internuclear distance in fused cells. Black dots represent cells in which both nuclei reside within the soma. The horizontal line denotes the distance threshold separating somatic from process-localized nuclei (n = 62). (H) Frequency distribution of internuclear distances. The histogram shows the number of fused cells per distance interval. The blue line shows the cumulative percentage of cells in each distance range.

After all the analysis, brightness and contrast were adjusted in all images for figure preparation.

### Statistical analysis

Statistical analysis was performed using GraphPad Prism 9. Paired analysis was performed to compare two samples. Student’s paired t-test was used for data that passed the normality test, whereas the Wilcoxon test was used when the normality test failed. Multiple sample comparisons were performed using one-way ANOVA and Šídák’s multiple comparisons test for normally distributed data. Data that failed the normality test were analyzed using either the Friedman test for paired data or the Kruskal-Wallis test for non-paired data, with the subsequent Dunn’s multiple comparisons test. Graphs display the mean when data pass the normality test and the median otherwise. Estimation plots display paired data points connected by lines in the left panel and the mean difference with 95% confidence interval (CI) in the right panel.

PCA was performed on the eight parameters described above after standardization (mean = 0, standard deviation = 1 across features). The two principal components (PCs) explaining the most variance (i.e., with the largest eigenvalues) were retained.

## RESULTS

### Mild EAE boosts infiltration and fusion of BMDCs in the cerebellum

BMDC infiltration and fusion with Purkinje cells occur at an extremely low rate under healthy conditions. Thus, we decided to study them in the neuroinflammatory context of EAE, where both events are substantially enhanced ^4,13^. To balance adequate cellular infiltration and fusion events with minimal neurological compromise, we optimized a mild EAE induction protocol following bone marrow transplantation (Fig. 1A). To do so, we first tested the effects of administering different amounts of MOG_35-55_ (75, 100, or 200 µg) to induce EAE. We did not observe any relevant sex-related difference or any difference between the 75 μg and 100 μg doses, but the maximum clinical score was significantly higher with the 200 μg dose (Fig. 1B). Besides, only 50% of the mice treated developed symptoms when EAE was induced with the 75 μg and 100 μg doses (4/8 mice per group, Fig. 1C), whereas all animals injected with 200 μg of MOG_35-55_ developed symptoms. Additionally, with the 200 μg dose, only 4 of 14 animals reached a maximum clinical score greater than 2, and no mice died from EAE or were sacrificed due to severe suffering (neither during this optimization study nor in subsequent analyses). These findings validate the successful establishment of a mild EAE model in which fusion events can be studied while mice exhibit minimal pathological changes. Therefore, we decided to use the 200 μg dose for the rest of our study.

In parallel, we wanted to explore infiltration and fusion at earlier EAE stages than previously reported, to increase the probability of capturing new, relevant biological information. Thus, we reduced the time window between bone marrow transplantation, EAE induction, and sacrifice compared with that previously used ^4,13^. We induced EAE one month after bone marrow transplantation, giving the mice just enough time to recover. We decided to euthanize the animals at either 8 or 14 weeks after EAE induction (Fig. 1A), when they were 15 and 21 weeks old, respectively, corresponding to approximately one-third and one-half of the age used in previous studies ^4,13^. In both cases, animals exhibited a typical EAE course, with a variably delayed peak in clinical score (some animals developed a multiphasic EAE) and full or partial recovery. For both sacrifice time points, we detected higher cell infiltration compared to control mice (Fig. 1D). Most GFP^+^ infiltrated cells expressed Iba1, a canonical marker for macrophage/microglial cells (Fig. 1E). As previously described, we also observed GFP^+^ heterokaryon Purkinje cells (Fig. 1F) clearly showing two different nuclei (arrow and arrowhead in Fig. 1F), as a consequence of cell fusion. Since we detected infiltrated and fused cells in sufficient numbers for our purposes at both time points, we decided to sacrifice mice at the earliest one (8 weeks after EAE) for the rest of our study.

In conclusion, we optimized a protocol that induced mild EAE, with few animals reaching a clinical score of 3 and none reaching the humane endpoint. This protocol enabled us to study cell infiltration and fusion approximately 15 weeks after birth (8 weeks after EAE induction), yielding a substantial number of fusion events and substantially reducing the time required in previous studies.

### Identification of BMDCs with BG identity

Our first step was to characterize fused and infiltrated cells in our EAE model. Among them, we identified some GFP+ cells exhibiting features distinct from previously characterized infiltrating or fused cell types. (Fig. 2A). They showed a small soma in the Purkinje cell layer, lying between Purkinje cells, and long processes displayed along the molecular layer, towards the pia mater (Fig. 2B). The position and morphology of these GFP^+^ cells were typical of BG, as shown by comparison with control BG detected by their expression of S100β and the glial fibrillary acidic protein (GFAP) markers ^43^ (Fig. 2C-D). They also showed short thorn processes in their fibers, typical of BG ^44^. We confirmed that these GFP^+^ cells were indeed BG by their expression of the typical BG markers GFAP and S100β in both their soma (Fig. 2E) and their processes (Fig. 2F). Furthermore, their staining pattern was indistinguishable from that of non-GFP neighbor BG cells. Thus, both morphology and marker expression confirmed that some of the GFP^+^ cells we observed in EAE-induced mice were indeed GFP^+^ BG.

### Bone marrow-related BG emergence is independent of EAE severity

We studied the distribution of GFP^+^ BG in the cerebellum of 14 EAE-induced and 3 control mice. Analyzing two out of six series of sections collected from half of the cerebellum (half vermis plus one hemisphere), we found a total of 1 to 12 cells per mouse (colored symbols in Fig. 3A). Thus, we estimate that the total number of BM- derived BG per EAE mouse cerebellum may range between 6 and 72 (1-12 x 6/2 series x 2 halves). However, we did not find any GFP^+^ BG in control animals (n = 3). To explore whether the emergence of these cells was related to the clinical score, we used the normalized cumulative disease index (CDI, see methods). When we included control animals in our correlation analysis, we observed a significant correlation between the normalized CDI and the number of GFP^+^ BG cells observed (p = 0.0274, r = 0.5396). Nevertheless, when control animals were excluded from the analysis, the correlation lost significance (p = 0.4396, r = 0.2235). Indeed, in addition to the 3 control animals, we did not find any GFP^+^ BG in two (out of 14) of all the EAE mice analyzed (white dots in Fig. 3A), although they reached maximum clinical scores of 0.5 and 2.5, and their normalized CDIs were 0.025 and 0.42. Furthermore, we did not observe a significant correlation between the animal’s weight loss, a common symptom of EAE mice ^45^, and the number of GFP^+^ BG.

Our data suggest that EAE-associated pathology is required but not sufficient for the emergence of bone marrow-related BG cells, and that the clinical severity of the EAE does not correlate with the abundance of these cells.

### GFP^+^ BG are heterogeneously distributed within the cerebellum of EAE mice

To deeply explore GFP^+^ BG distribution, we charted the locations of these cells (colored symbols in Fig. 3A) in each slice. We then grouped the sections into three major anatomical regions (Fig. 3B1-3), based on their lateromedial position (see methods). Only one GFP^+^ BG was found in the flocculus (upper left sections in Fig. 3B), which was not considered for this part of the study. Furthermore, within the vermis, we also analyzed the distribution of cells by local region (nodular, posterior, central, and anterior; Fig. B3, D) and by cerebellar lobe (as in the vermis, all lobes can be unequivocally distinguished; Fig. B3, E). Cell distribution was heterogeneous at different levels. First, the number of cells we found in the vermis (36 in total) doubled the number of cells found in the cerebellar hemispheres (18 in total; p = 0.0020; Fig. 3C). We thought that this could be because the length of the Purkinje cell layer was apparently larger in the vermis than in the hemisphere. To evaluate this possibility, we measured the length of the Purkinje cell layer in each section and calculated the linear density of GFP^+^ BG cells. Still, the linear density was significantly higher in the vermis than in the hemispheres, suggesting that GFP^+^ BG arise with higher frequency in the vermis (p = 0.0024; Supp. Fig 1).

Within the vermis, the anterior domain was the region where GFP^+^ BG cells were more abundant. In particular, the number of cells in the anterior domain was significantly higher than in the posterior (p = 0.0339) and nodular domains (p = 0.0017; Fig. 3D). Finally, we studied the distribution within vermal lobes. Interestingly, lobes IV-V and VI accounted for more than 60% of the total number of positive cells, while we did not find any GFP^+^ BG in lobe X (Fig. 3E).

Altogether, these results demonstrate that GFP^+^ BG are distributed heterogeneously in the cerebellum, with their predominant location in lobes IV-V and VI of the vermis.

### GFP^+^ BG are the result of the fusion between BMDCs and BG

The presence of GFP^+^ BG could be explained by either BMDC transdifferentiation or fusion with resident BG. The main structural difference between these two mechanisms is that cell fusion leads to the formation of heterokaryons, that is to say, cells with two different nuclei ^3^. Thus, we used a combination of regular confocal imaging and super- resolution (STED) imaging to check whether GFP^+^ BG had one or two nuclei. Among the 80 analyzed GFP^+^ BG cells, we identified 63 cells with two nuclei from 11 different animals. Of these, 38 cells showed two nuclei in their soma. We confirmed that both nuclei (Fig. 4A, arrow and arrowhead; supplementary videos 1 and 2) were surrounded by GFP and GFAP staining, demonstrating that they belonged to the same BG cell. Importantly, the cells in which we were not able to identify two nuclei were not fully included in the cerebellar section; thus, we cannot exclude the presence of two nuclei in them.

We also observed many GFP^+^ BG cells with only one nucleus in the soma, so we took a deeper look at their processes and, surprisingly, discovered 25 additional cells with a second nucleus in their processes. Some of these cells had an additional nucleus very close to the soma, whereas in other cases the nucleus was further away, either towards the granule cell layer (Fig. 4B) or to the molecular layer (Fig. 4C-D). We used 3D reconstructions to confirm that the displaced nucleus (Fig. 4B-C, arrowhead) was structurally connected via cellular processes to the GFP^+^ BG soma (Fig. 4B, dotted arrow) and fully enclosed by the GFP^+^ cytoplasm (Fig. 4C). In all these analyzed heterokaryons, we confirmed the existence of a second nucleus (Fig. 4B-C, arrow) within the soma of the BG cell, which should be the original nucleus of BG cells. Furthermore, super-resolution STED microscopy of DAPI, GFP, and Lamin B2 staining confirmed that displaced nuclei were strictly contained within the GFP^+^ boundaries of the BG cells (Fig. 4E). In total, we detected 63 fused cells out of 80 GFP^+^ BG cells, suggesting that cell fusion is a potent and widespread phenomenon in BG. The remaining 17 cells were excluded from the analysis due to their superficial location within the cerebellar section, which prevented full volumetric assessment to confirm or rule out the presence of two nuclei.

To characterize the spatial integration of the donor nuclei, we mapped their position relative to the original BG nucleus. We observed that both nuclei co-localized within the soma in approximately 60% of the cases (Fig. 4F). To further resolve the spatial distribution of the non-somatic nuclei, we measured the 3D distance between the centroids of both nuclei using z-stack images. Inter-nuclear distances ranged from ∼2 μm to 88 μm. The frequency distribution of these distances, plotted in Figure 4G, revealed a clear skew toward shorter values, indicating that BMDC nuclei are preferentially located in the proximal regions of the processes rather than at the distal apical tips (Fig. 4H). This centripetal distribution suggests that, although fusion may occur in the processes, the BMDC nucleus tends to be positioned in or near the soma.

Altogether, our observations demonstrate that GFP^+^ BG cells result from the fusion of a BMDC and a BG, characterized by a preferential nuclear distribution where the donor nucleus is positioned within or near the host soma.

### Nuclei in fused BG exhibit distinct morphological and chromatin activation profiles

Once we confirmed that GFP^+^ BG resulted from fusion between BG and BMDCs, we next characterized the differences between the two nuclei of each cell using DAPI staining to delineate nuclear boundaries and analyze nuclear characteristics. We reasoned that the nuclei observed in the processes of GFP^+^ BG cells, which were consistently small and round, were bone marrow-derived (Figure 4B-D, arrowhead). Importantly, when both nuclei were in the soma, one was consistently smaller with a more rounded contour than the other, resembling those found in the processes. Therefore, we classified this distinct nucleus, which could be in the soma or in the processes, as the BMDC-derived one (Fig. 5A). The morphological differences observed between nuclei support the notion that fused BG are heterokaryons, consistent with previous reports of bone marrow-fused Purkinje cells exhibiting nuclear heterogeneity ^3^.

**Figure 5.**
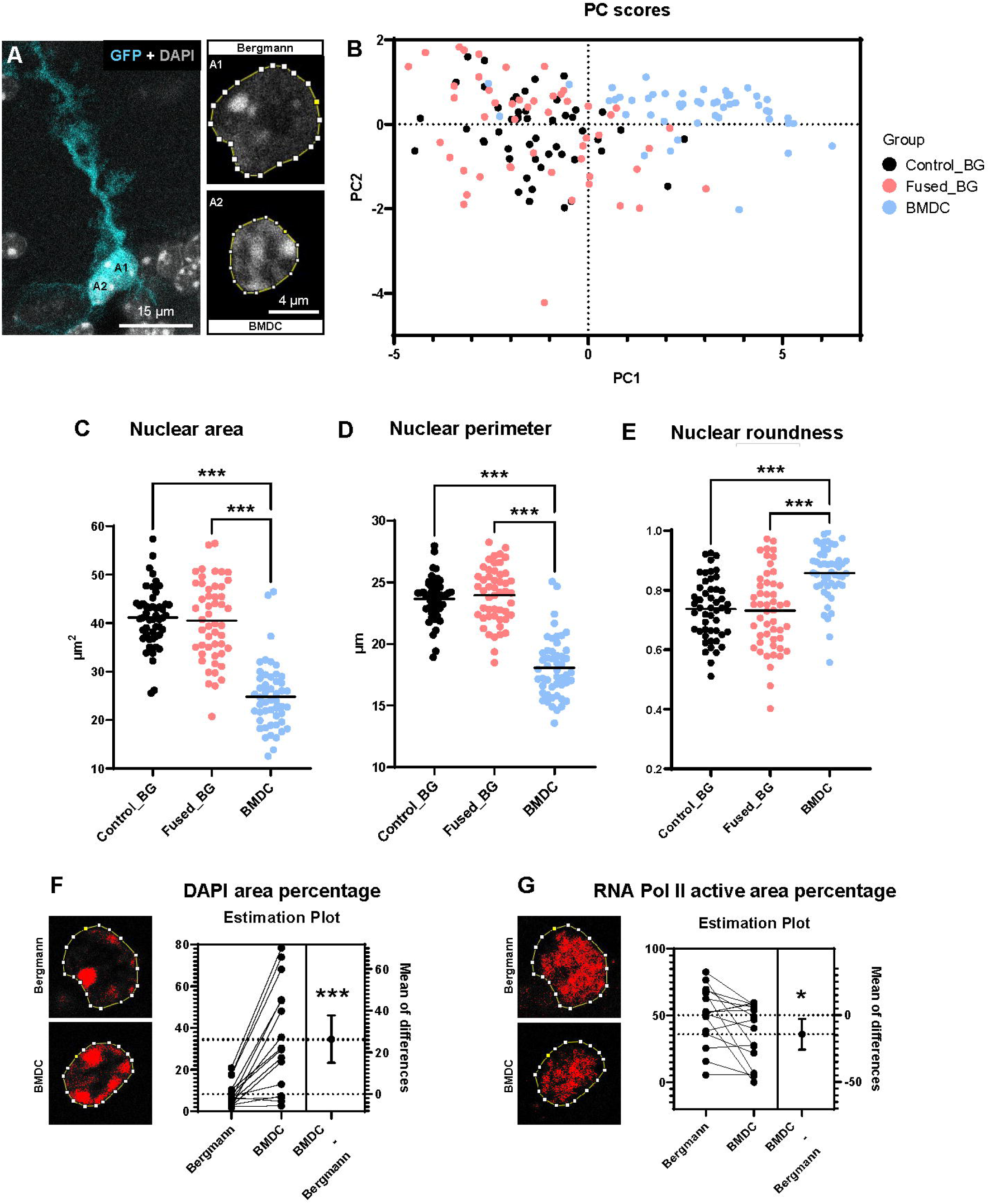
Morphological and functional differences between donor and resident nuclei within fused BG. (A) Representative image of fused, GFP⁺ (cyan), BG showing ROIs created for the segmentation of the two different nuclei based on DAPI staining (gray). A1 shows the original nucleus from BG, while A2 shows the BMDC nucleus. (B) PCA of 150 nuclei. The plot reveals distinct clustering of nuclei classified as BMDC nuclei (cyan) and those classified as resident BG nuclei within heterokaryons (orange). The latest clusters with nuclei from control BG (black). (C–E) Quantitative comparison of nuclear area (C), perimeter (D), and roundness (E) for control BG nuclei (black), original BG nuclei from fused cells (orange), and BMDC nuclei from fused cells (cyan). Solid lines indicate the mean (one-way ANOVA with Šídák’s multiple comparisons test; n = 50 nuclei per group). (F) Analysis of chromatin condensation. Left: images of the nuclei illustrating the threshold used for DAPI area percentage analysis. Right: estimation plot showing the paired relationship between original BG nuclei and BMDC nuclei within the same fused cells (paired t-test; n = 16 nuclei per group). (G) Analysis of transcriptional activity. Left: images of the nuclei illustrating the threshold used for active RNA polymerase II area percentage analysis. Right: estimation plot showing the paired relationship between original BG nuclei and BMDC nuclei from fused cells (paired t-test; n = 16 nuclei per group). * p < 0.05; *** p < 0.001.

To assess the validity of our classification, we analyzed the morphology of control, non-fused BG nuclei and compared it with the morphology of GFP^+^ heterokaryon BG nuclei. Nuclei located within the processes served as a reference for BMDC nuclei with a clear identity. We selected eight morphological parameters quantified by FIJI (area, perimeter, width, height, major and minor axis lengths of the best fitting ellipse, Feret’s diameter or maximum caliper, and minimum Feret’s diameter or minimum caliper) for PCA. Derived shape descriptors (e.g., circularity, which reflects the ratio of area and perimeter) were excluded to avoid redundancy. PC1 explained 80.14% of the total variability of our data, and together with PC2, they accounted for 91.28% of the variance. When we plotted the PC1 and PC2 scores for each nucleus, most of those classified as bone marrow-derived clustered in a distinct region of the graph, separated from nuclei identified as BG-putative in heterokaryons. In contrast, most reference nuclei from control BG, and nuclei classified as the original from BG in GFP^+^ cells appeared clustered together in a separate region of the plot (Fig. 5B). These results demonstrated the validity of our visual classification criteria.

To further validate our classification strategy and the morphological differences observed between nuclei of different origins, we performed a correlation matrix analysis of the eight morphological variables used within the PCA. By comparing each pair of morphological variables (Suppl Fig. 2), we found that most of them clearly captured differences between nuclei derived from BMDC and those from the original BG. As described for the PCA, in most correlation graphs, BMDC nuclei segregated from BG- nuclei (either from heterokaryon GFP-positive cells or from non-fused GFP-negative control cells). Six of the 28 analyzed correlations showed a Pearson’s correlation coefficient above 0.9: area vs perimeter, major axis vs perimeter, Feret’s diameter vs perimeter, Feret’s diameter vs major axis, minimum Feret’s diameter vs major axis, and minimum Feret’s diameter vs minor axis. The area and perimeter showed the highest correlations with the other variables (r = 0.777-0.967 for the area and r = 0.722-0.967 for the perimeter), as expected given their relevance to our cell classification method based on nuclear size. Notably, the perimeter and major axis variables were the most frequent (3 vs 2 for Feret’s related variables, and vs 1 for area and minor axis) for correlations with an r higher than 0.9. Indeed, when each parameter was plotted against itself, BMDC nuclei were clearly differentiated, in most cases, from fused and control BG nuclei (Suppl Fig. 2), showing that one of the eight parameters could suffice to classify the nuclei.

To statistically demonstrate the differences observed in BMDC and BG nuclei size and shape, we analyzed their area, perimeter, and the ImageJ shape descriptor roundness (based on the relationship between the area and the major best-fitted ellipse axis), which reflects the elongation of the nuclei, ranging from 0 to 1 (1 corresponding to a perfect circle). We found no difference between control and BG-putative nuclei in heterokaryons across all parameters analyzed, further validating our visual classification. Nevertheless, the nuclei from the BMDC were significantly different from BG nuclei in size and shape. Their area and perimeter were smaller (p < 0.0001; Fig. 5C, D), and their roundness higher (p < 0.0001; Fig. 5E). Therefore, our results indicate that, in fused cells, the original BG nucleus does not alter its size or morphology to accommodate the incoming nuclei. In addition, BMDC nuclei are clearly distinct from BG nuclei, exhibiting a smaller and rounder profile, whereas BG nuclei display a more elongated, ellipsoid shape.

Finally, to link nuclear morphology with function, we studied the transcriptional activity of both nuclei within fused cells. We measured the percentage of nuclear area covered by highly intense DAPI regions and by the active form of RNA polymerase II as markers of heterochromatin and transcriptional activity, respectively. To avoid potential research bias, we analyzed all possible intensity thresholds, observing significant differences across a wide range of values (Suppl. Fig. 3). From these, we selected the threshold that best captured the heterochromatin patches visualized by DAPI staining and active transcriptional regions marked by the active form of RNA polymerase II immunostaining (Fig. 5F-G). The area percentage covered by intense DAPI staining was significantly higher in BMDC nuclei than in BG nuclei (p = 0.0002; Fig. 5F), indicating that BMDC nuclei are more heterochromatic than those of BG. Conversely, the percentage of area covered by the active form of RNA polymerase II was higher in Bergmann nuclei than in BMDC nuclei (p = 0.0179; Fig. 5G). These findings demonstrate that, within fused BG, nuclei of BMDC origin remain transcriptionally active, albeit at a lower level compared to the original BG nuclei.

Taken together, our quantitative analysis reveals that fused Bergmann glia are heterokaryons exhibiting pronounced nuclear asymmetry, characterized by the coexistence of two morphologically and functionally distinct nuclei. The persistence of a compact, less active BMDC nucleus alongside a large, transcriptionally active resident BG nucleus implies that the donor genetic material is integrated without disrupting the morphology and general transcriptional activity of the host nucleus.

## DISCUSSION

Here, we demonstrate that BMDCs infiltrate the brain and fuse with BG during EAE. We detected these fused cells 8 weeks after EAE induction, noting that their presence is independent of the severity of the EAE-related symptoms. Fused BG retain typical BG markers, and their distribution is heterogeneous within the cerebellum, preferentially populating the vermis. While we found most BMDC nuclei in BG soma, as typically described for Purkinje cell fusion, we also detected them within BG processes, suggesting that cell fusion may occur in this part of the cells. Morphologically, the resident BG nucleus remains unchanged (indistinguishable from those of non-fused BG), whereas the incoming BMDC nucleus is distinctively smaller and more circular than the original BG nucleus. Crucially, both nuclei remain transcriptionally active, although the BMDC nucleus displays lower transcriptional activity and greater heterochromatinization. Thus, we undoubtedly identify these fused BG cells as heterokaryons, as others have previously done with fused Purkinje cells ^3^.

Our initial objective was to establish an experimental protocol to study fusion between BMDCs and Purkinje cells in the EAE model, optimizing the balance between the number of detectable fusion events and a practical experimental timeframe. Fusion between BMDC and Purkinje cells was initially described in healthy mice following bone marrow transplantation ^1^, and subsequent studies demonstrated that the number of fused cells increases progressively with time ^3^. Later, an increase in cell fusion was reported under pathological conditions in both mice ^4,9,46^ and humans ^10,12^, with particularly pronounced effects observed in EAE mice ^4,13^. However, experiments in EAE mice have typically required approximately 10 months. In the present work, we shortened the overall timeline -encompassing birth, BMDC transplantation, EAE induction, and endpoint analysis- to approximately four months. This optimized protocol allowed us not only to detect Purkinje cell fusion (data not yet published) but also to identify BMDC fusion with BG. For future investigations, we propose this timeframe as a suitable window for quantitative analysis, while recognizing that additional post- induction intervals may be necessary to fully capture earlier initiation or long-term dynamics of these fusion events.

We have described, for the first time, fusion between BMDC and BG in the cerebellum of EAE mice. After the discovery of fusion between BMDCs and Purkinje cells, different authors have studied the ability of BMDCs to fuse with other cell types in the brain. In the context of EAE, fusion has also been observed with spinal cord motor neurons, forming heterokaryons ^23^. As well, fusion with neurons has been described in the retina after damage ^47^ and in the substantia nigra pars compacta of a mouse model of Parkinson’s disease ^24^. Surprisingly, Altarche-Xifro and collaborators also reported the presence of fused glial cells (GFAP^+^) in their study ^24^. Some years later, researchers from this same group demonstrated fusion between BMDCs and Müller Glia cells in an animal model of retinal degeneration induced by N-methyl-D-aspartate ^26^. Finally, Chen et al. (2011) reported sideways GFP^+^ BG cells following GFP^+^ BMDC transplantation while investigating Purkinje cell fusion in a transgenic model of spinocerebellar ataxia type 1. Although they did not confirm cell fusion and their paradigm was based on genetic neurodegeneration rather than autoimmunity, as in our EAE model, they detected such cells in 4 of 10 animals. Our findings definitively demonstrate that the fusogenic potential of BMDCs in the cerebellum is not limited to Purkinje cells but also encompasses BG.

Our data strongly support cell fusion as the main mechanism underlying GFP^+^ BG formation. Nevertheless, we wanted to discard other possible mechanisms. Beyond fusion, transdifferentiation is the other mechanism by which BMDCs can integrate into the brain^35^. Bone marrow stromal cells injected into neonatal brains can differentiate into GFAP⁺ astrocytes ^48^, and in adults, bone marrow–derived cells can give rise to both microglia and macroglia ^49^. However, BG generation via transdifferentiation or heterokaryon transdifferentiated cells has never been documented. A scenario in which a BMDC transdifferentiates and subsequently undergoes karyokinesis would initially generate mononucleated GFP⁺ BG cells, a phenomenon we did not observe. We also considered the option of BG phagocytosing BMDCs. Under certain circumstances, microglia can phagocyte non-apoptotic cells in a process known as “phagoptosis” ^50^. Astrocytes, and specifically BG, have phagocytic activity ^51,52^, and some authors suggest that, during microglial dysfunction, astrocytes may also exhibit phagoptosis ^53^. However, after phagoptosis, the membrane of the engulfed cell remains intact ^54^ such that GFP expression would be confined to the internalized cell (a phenomenon we never saw), unlike the homogeneous GFP labeling we observed throughout the BG cytoplasm. These results reinforce cell fusion as the origin of GFP^+^ BG, rather than other mechanisms.

Fused BG cells exhibited a heterogeneous distribution across the cerebellum, predominantly localizing to the vermis as previously reported for fused Purkinje cells ^23^. This pattern likely reflects the vermis’s heightened vulnerability to age-related degeneration ^55^. Notably, this susceptibility extends to finer lobe-level differences, as we detected no fused cells in lobe X, which is a known neuro-resistant region ^56–58^. Furthermore, it has been demonstrated that BG cells are organized into topographically distinct molecular zones with region-specific gene expression profiles ^59^, which could also modulate their propensity to undergo fusion. Future studies should dissect how these molecular gradients, together with damage-associated signals, regulate fusion likelihood across cerebellar domains.

Most fused cells contained both nuclei within the soma, but some exhibited one nucleus positioned along their processes, suggesting that fusion may occur not only in the soma (as previously reported for Purkinje cells^1–4,9,13,18^) but also along the processes. Furthermore, within the processes, most nuclei were close to the soma, which led us to hypothesize that they might be migrating towards the soma, where both nuclei are ultimately found in most fused BG cells. Previous studies of cell fusion in the brain consistently reported that both nuclei reside within the soma ^1,4,13,23,24^, suggesting that this localization represents the stable configuration. Concentrating both nuclei within the soma, where most organelles reside, would also constitute the most energy- efficient arrangement. Analogously, during myogenesis, fused myoblasts form multinucleated myotubes in which nuclei actively translocate and cluster at central positions ^60^. Likewise, during development, the primary BG nucleus migrates from its process into the soma ^61^. Together, these observations suggest that nuclear migration toward the soma may be a conserved feature of post-fusion remodeling. However, additional studies are required to determine whether BMDC nuclei invariably relocate to the soma, persist stably within the process, or instead migrate from the soma to the processes.

Collectively, our findings reveal an unanticipated dimension of bone marrow–derived cell plasticity within the cerebellum under pathological conditions. Beyond the previously recognized fusion with Purkinje neurons, we demonstrate that BG can also form stable heterokaryons with infiltrating BMDCs, preserving glial identity while accommodating a second transcriptionally active nucleus. The discovery that fusion may occur not only in the soma but also along BG processes supports a new mechanistic model for BMDC–BG interaction. Moreover, the marked morphological and transcriptional asymmetry between the two nuclei underscores functional heterogeneity that may influence glial physiology and cerebellar homeostasis. These findings strikingly expand the conceptual framework of cell fusion in the CNS, positioning BG as a previously unrecognized partner in neuroinflammatory remodeling and highlighting fusion-based strategies for neuroregeneration.

## AUTHOR CONTRIBUTION

P.G.TdM., D.D., J.V., and E.W. conceived the project and designed the study. P.G.TdM. performed the EAE protocol. P.G.TdM., S.S-M. and C.E-S performed histological procedures and analyzed the data. P.G.TdM. and C.A.A-Z performed the statistical analysis. E.W., D.D., J.V., obtained the funding. P.G.TdM. wrote the manuscript and prepared the figures. D.D., J.V., and J.R.A reviewed and edited the manuscript. D.D., J.V., and J.R.A supervised the scientific work. J.V and J.R.A contributed equally to this work and share last authorship. All the coauthors critically revised and approved the final manuscript.

## ACKNOWLEDGMENTS

We thank the CIC-IBMCC Advanced Cellular Analyses Unit for technical assistance with STED imaging and Luis Antonio de Castro for support with confocal imaging. We are also grateful to Javier Borrajo, from the Radioisotopes and Radiation Protection Unit, Nucleus Service, Universidad de Salamanca, for performing mouse irradiation. P.G.TdM is funded by the Spanish Ministry of Universities (FPU20/03457). The whole project was funded by the Spanish Ministry of Science and Innovation (PID2022- 140525NB-I00, PID2022-140456NB-I00, PID2019-106943RB-100), the Strategic research programmes of excellence from JCyL, co-funded by the ERDF Operational Programme (CLU-2023-1-01) the JCyL (SA112P24, SA129P20) and by the Universidad de Salamanca.

## FIGURES

**Supplementary Figure 1. Regional comparison of GFP**^⁺^ **BG density.** Quantification of the linear density of GFP⁺ BG cells comparing cerebellar sections from the vermis versus non-vermal regions. Solid line indicates the median (n = 12; Wilcoxon test). **p < 0.01.

**Supplementary Figure 2. Correlation matrix of nuclear morphological parameters.** Scatter plot matrix showing the pairwise relationships between the eight shape descriptors utilized for PCA. Pearson correlation coefficients (r) are shown for each comparison. The diagonal panels display each parameter plotted against itself. Black dots represent nuclei from control BG, orange dots represent original BG nuclei from fused cells, and cyan dots represent donor BMDC nuclei.

**Supplementary Figure 3. Staining analysis with Thresholderer**. Curves showing the percentage of nuclear area covered by DAPI staining (A) or the active form of RNA polymerase II staining (B) as a function of increasing threshold values. Solid colored lines indicate the mean, and dotted lines with shaded regions represent the standard error of the mean.

