## Supplementary figures and images for "Bone Marrow-Derived Cells Fuse with Bergmann Glia in Experimental Autoimmune Encephalomyelitis"

### Supplementary figure 1

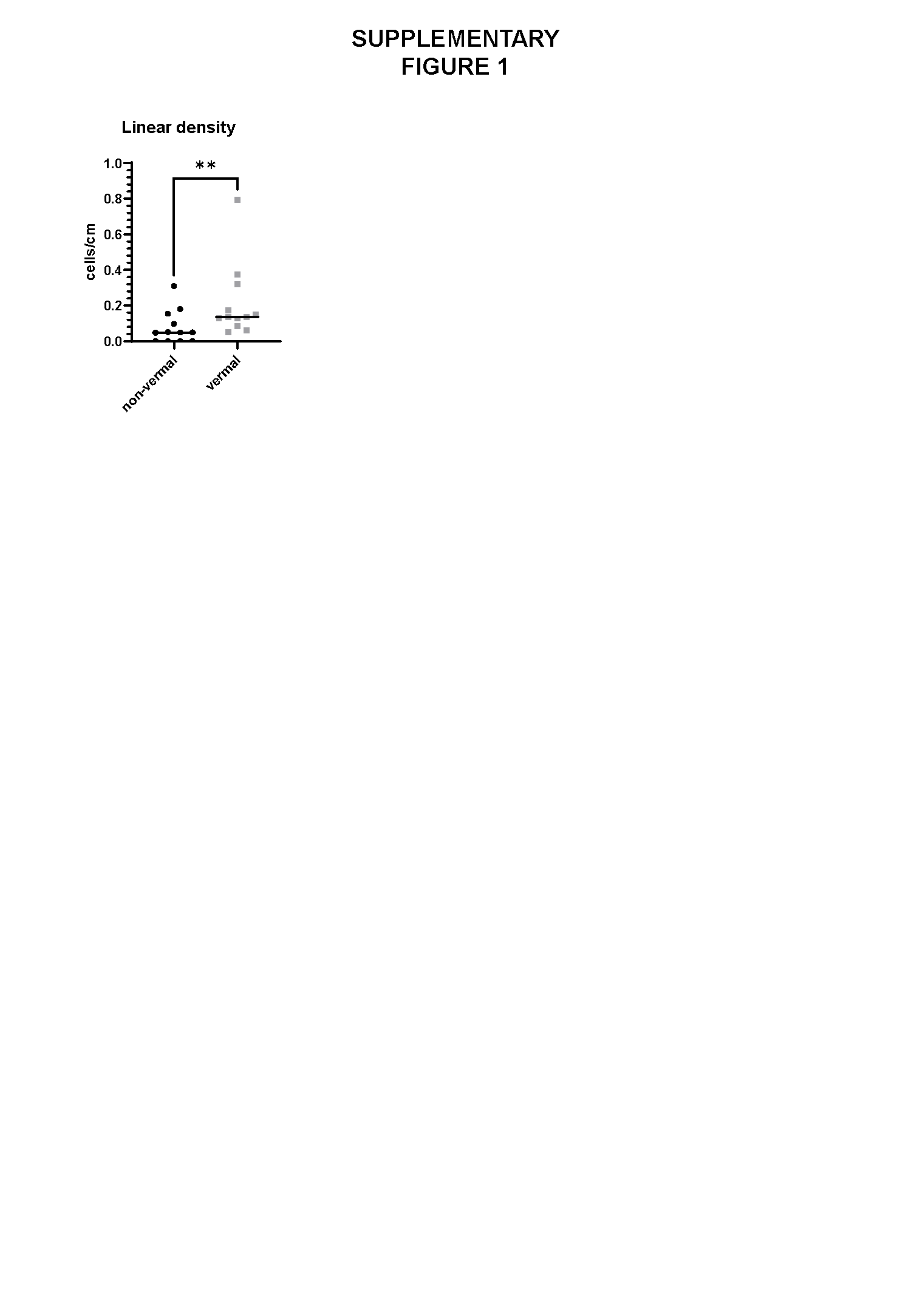

### Supplementary figure 2

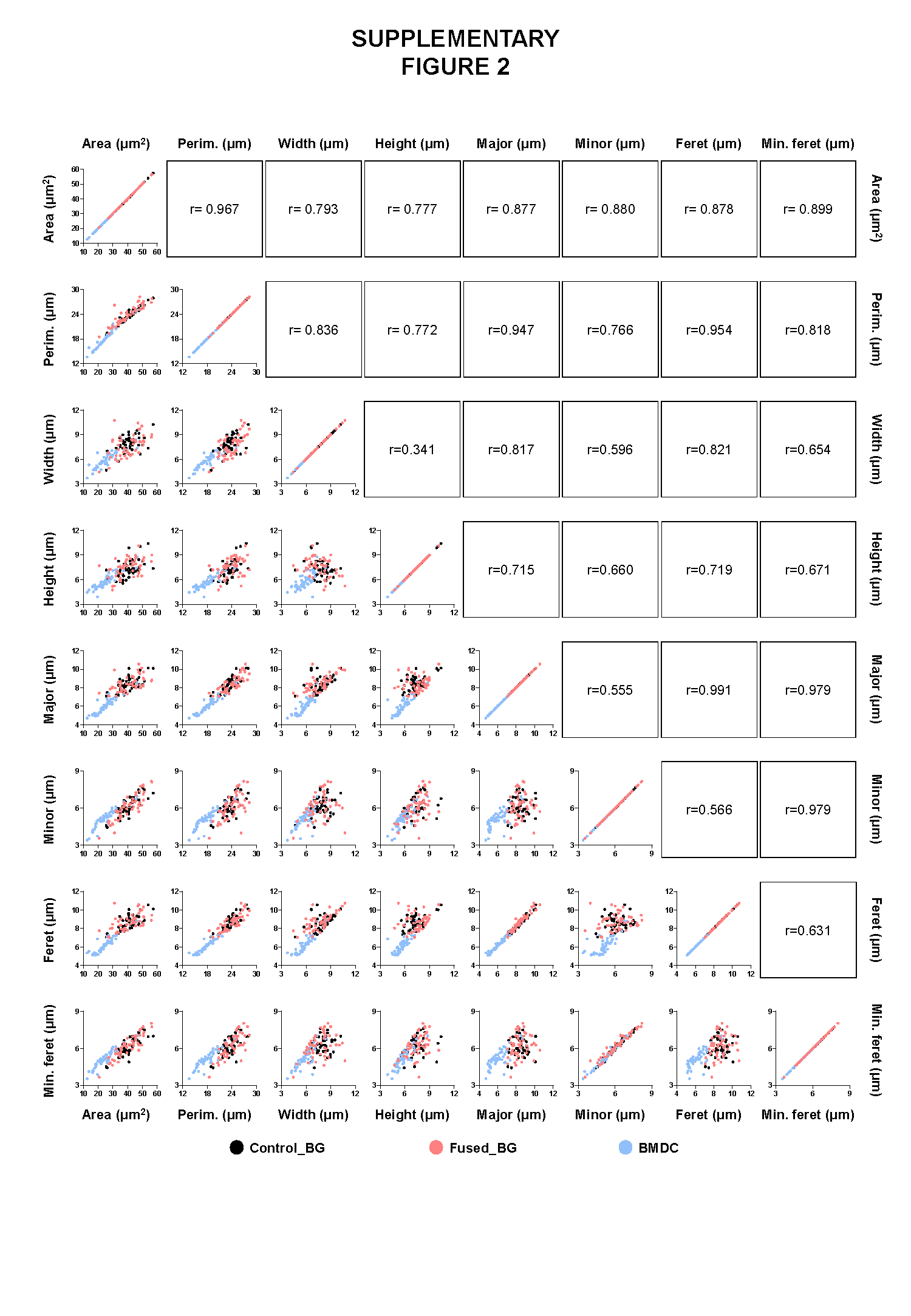

### Supplementary figure 3

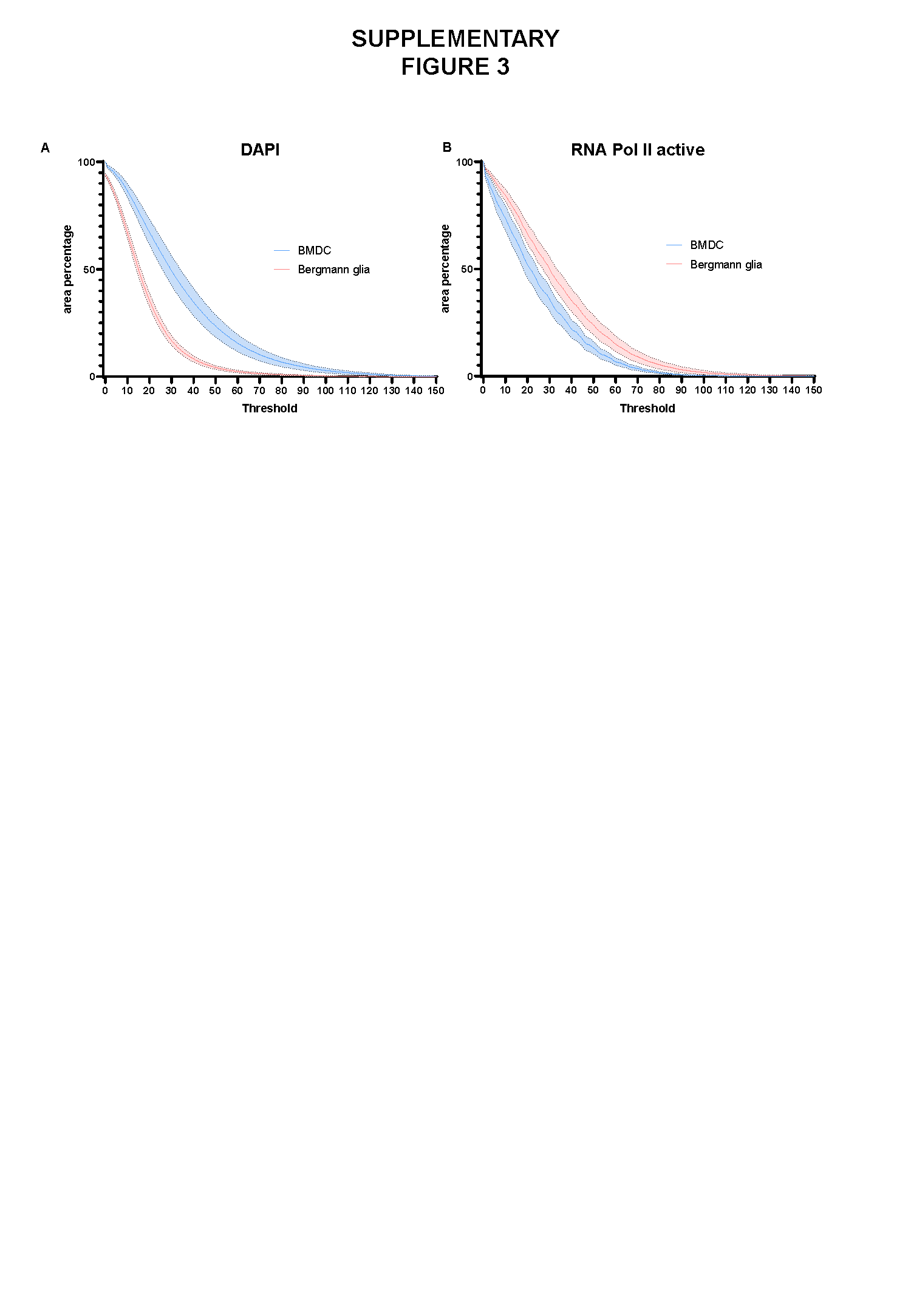
