## Supplementary materials for "Bone Marrow-Derived Cells Fuse with Bergmann Glia in Experimental Autoimmune Encephalomyelitis"

### BG linear density estimation

Mosaic images were used for the linear density analysis. For each cerebellar section, the length of the Purkinje cell layer was measured in FIJI using the segmented line tool, and the number of GFP+ Bergmann glia was divided by the cumulative Purkinje layer length of either vermal or non-vermal sections to obtain a linear cell density (cells per centimeter).

### Correlation matrix statistics

A correlation matrix was computed using eight nuclear shape descriptors: area, perimeter, width, height, major and minor axis lengths of the best-fitting ellipse, Feret's diameter or maximum caliper, and minimum Feret's diameter or minimum caliper. All pairwise comparisons were analyzed. The resulting matrix was visualized as a scatterplot–correlogram, where the lower triangle showed the bivariate scatter plots for each parameter pair, the upper triangle reported the corresponding Pearson correlation coefficients, and the diagonal panels displayed each parameter against itself to visualize group-wise distributions.

### Fused Bergmann glia video

GFP-positive signals were segmented as described in the Methods section "*Data collection and image analysis.*" The video was generated using the "3D Project" tool in FIJI, applied to the full z-stack dataset. The projection method was set to *Brightest Point*, with rotation around the Y-axis. All other parameters were kept at default, and the *Interpolate* option was enabled. Brightness and contrast were adjusted for optimal visualization. The final video was exported in AVI format (uncompressed), at a resolution of 1624 × 1568 pixels and 10 frames per second.
