## Supplementary material for "Bone Marrow-Derived Cells Fuse with Bergmann Glia in Experimental Autoimmune Encephalomyelitis": Table 1

| Antibody | Host Species | Reference | Clone / Lot | Dilution | Supplier |
|---|---|---|---|---|---|
| Anti-Calbindin-D28k | Mouse | AB_10000347 | 07 | 1/2000 | Swant |
| Anti-GFP | Goat | Ab5450 | GR3371088-1 | 1/2000 | Abcam |
| Anti-GFP | Mouse | Ab291-50 | 383290 | 1/1000 | Abcam |
| Anti-Lamin B2 | Rabbit | AB_2802716 | WI3376410 | 1/1500 | Invitrogen |
| Anti-GFAP | Mouse | G6171 | 082M4808 | 1/1000 | Sigma-Aldrich |
| Anti-S100β | Rabbit | Ab5264 | 1068686-30 | 1/4000 | Abcam |
| Anti-RNA Polymerase II | Rabbit | Ab5095 | 1076049-2 | 1/2000 | Abcam |
| Anti-Iba1 | Goat | Ab5076 | GR153106-4 | 1/1000 | Abcam |
| Cy2 anti-goat | Donkey | 705-225-147 | 133888 | 1/500 | Jackson ImmunoResearch |
| Cy2 anti-mouse | Donkey | 715-225-150 | 100331 | 1/500 | Jackson ImmunoResearch |
| Cy3 anti-mouse | Donkey | 715-165-150 | 122771 | 1/500 | Jackson ImmunoResearch |
| Cy3 anti-goat | Donkey | 705-165-147 | 79773 | 1/500 | Jackson ImmunoResearch |
| Cy5 anti-rabbit | Donkey | 711-175-152 | 84963 | 1/500 | Jackson ImmunoResearch |
| Alexa 488 anti-goat | Donkey | AB_2534102 | 2059218 | 1/500 | Invitrogen |
| Alexa 555 anti-rabbit | Donkey | AB_162543 | 2482963 | 1/500 | Invitrogen |
